# mTORC1 is required for differentiation of germline stem cells in the *Drosophila melanogaster* testis

**DOI:** 10.1101/2022.07.25.501357

**Authors:** Marie Clémot, Cecilia D’Alterio, Alexa Kwang, D. Leanne Jones

## Abstract

Metabolism participates in the control of stem cell function and subsequent maintenance of tissue homeostasis. How this is achieved in the context of adult stem cell niches in coordination with other local and intrinsic signaling cues is not completely understood. The Target of Rapamycin (TOR) pathway is a master regulator of metabolism and plays essential roles in stem cell maintenance and differentiation. We observe differential expression of the Tor kinase in the *Drosophila* male germline, which correlates with restriction of mTORC1 activity to germline stem cells (GSCs) and early germ cells. Targeted RNAi-mediated downregulation of Tor in early germ cells causes a block and/or a delay in differentiation, resulting in an accumulation of germ cells with GSC-like features. These early germ cells also contain unusually large and dysfunctional autolysosomes. In addition, downregulation of Tor in adult male GSCs and early germ cells causes non-autonomous activation of mTORC1 in neighboring cyst cells, which correlates with a disruption in the coordination of germline and somatic differentiation. Our study identifies a previously uncharacterized role of the TOR pathway in regulating male germline differentiation.

## Introduction

Adult stem cells have the unique ability to both self-renew and produce daughter cells that acquire a distinct identity through differentiation, thereby playing essential roles in the maintenance of tissue homeostasis. A myriad of factors has been shown to contribute to the regulation of stem cell behavior, which include cell-intrinsic factors and extrinsic cues, either systemic or provided by the local micro-environment (or “niche”) [1–3]. In particular, the capacity of stem cells to self-renew or differentiate can be attributed to distinct metabolic states and metabolism has emerged as an important regulator of stem cell fate decisions [4–8]. Yet, how metabolic cues are integrated with other signaling pathways to influence stem cell behavior in the niche is not fully understood.

The testis of *Drosophila melanogaster* is an excellent system for investigating the mechanisms that regulate stem cell behavior. Furthermore, as it contains two populations of adult stem cells - germline stem cells (GSCs) and somatic cyst stem cells (CySCs), the testis provides an ideal model to address questions related to how multiple stem cell systems can be coordinated to maintain tissue homeostasis [9,10]. Recently, a screen conducted in our laboratory designed to uncover the role of mitochondrial dynamics in regulating *Drosophila* male GSC behavior showed that the mitochondrial fusion protein Mitofusin (dMfn) is essential for GSC maintenance [11]. Depletion of dMfn in early germ cells resulted in ectopic phosphorylation of 4E-BP, a substrate of Tor kinase, suggesting that the Target of Rapamycin (TOR) pathway was hyperactivated in these cells in the absence of dMfn. Consistent with this observation, the TOR inhibitor rapamycin partially rescued GSC loss induced by dMfn depletion in germ cells [11]. Therefore, TOR hyperactivity appears to contribute to GSC loss in the *Drosophila* testis upon perturbation of mitochondrial fusion in germ cells, raising the question of the role of TOR in the regulation of adult male GSCs under homeostatic conditions.

The TOR pathway is a highly conserved signaling network that functions as a metabolic rheostat, coordinating cellular growth and proliferation with the availability of resources [12–14]. The Ser/Thr kinase Tor is the catalytic core of the pathway and forms part of two distinct complexes: mechanistic target of rapamycin complex 1 (mTORC1) and mechanistic target of rapamycin complex 2 (mTORC2), which differ in accessory proteins, upstream regulatory signals and substrate specificity. The mTORC1 complex controls metabolic pathways in response to intracellular nutrients and energy levels, as well as extracellular factors such as hormones and growth factors, thereby integrating local and systemic cues. Upon activation, mTORC1 promotes anabolic processes, including protein, lipid and nucleotide synthesis, while repressing catabolic processes such as autophagy, whereas mTORC2 regulates survival and proliferation [15]. Given its central role in metabolism, the TOR pathway provides a tool to gain insights into how metabolic inputs are integrated with other signaling pathways to regulate stem cell fate decisions.

Indeed, mTORC1 regulates stem cell behavior and fate decisions across many systems, including murine spermatogonial progenitor cells, mesenchymal stem cells, hematopoietic stem cells, intestinal stem cells, and hair follicle stem cells, among others [16]. Interestingly, mTORC1 appears to be required for both stem cell maintenance and the differentiation of progenitor cells, suggesting that mTORC1 activity must be fine-tuned and tightly regulated [16].

The genetic toolkit available in *Drosophila* facilitates modulation of the TOR pathway in a temporal and tissue-specific manner, to probe the role of TOR in adult stem cell maintenance, rather than during development. For example, in the *Drosophila* ovary, TOR was found to be required for GSC maintenance and proliferation, as well as for the growth and survival of differentiating progeny [17]. By contrast, hyperactivation of mTORC1 through cell-specific disruption of its repressor Tuberous sclerosis complex (TSC1/2), causes precocious GSC differentiation [18]. In addition, precisely tuned mTORC1 activity controls the number of mitotic divisions and the mitotic to meiotic transition in germline cysts [19,20]. In the *Drosophila* testis, a role for mTORC1 in cyst cell differentiation has been described [21]; however, the role of TOR in adult male GSCs under homeostatic conditions has not been addressed.

Here, we find that differential expression of the Tor kinase in the male germline correlates with mTORC1 activity in GSCs and early germ cells. Consequently, depletion of Tor in early germ cells leads to a block or delay in differentiation and an accumulation of germ cells with GSC-like features. Germ cells depleted for Tor kinase activity also harbor unusually large and dysfunctional autolysosomes. Remarkably, downregulation of Tor in adult GSCs and early germ cells causes non-autonomous activation of mTORC1 in neighboring cyst cells, which correlates with precocious differentiation. Therefore, our study identifies a previously uncharacterized role of the TOR pathway in male germ cells, expanding what is known about this crucial regulator of metabolic and tissue homeostasis.

## Results

### Differential expression of Tor correlates with variations in TOR activity throughout germ cell differentiation

As a first step to investigate the role of TOR signaling in the adult male germline, we assessed the expression pattern of Tor, the core kinase of the pathway. The adult *Drosophila* testis is a coiled tube, in which spermatogenesis proceeds in a well-defined spatiotemporal manner. The stem cell niche is located at the apical tip and contains six to nine GSCs, surrounding a group of somatic support cells referred to as the hub. The hub is a critical signaling center that coordinates the maintenance of both GSCs and CySCs. GSCs divide asymmetrically to self-renew and give rise to a gonialblast (GB). GBs are displaced away from the hub and undergo four rounds of synchronous transit amplification (TA) divisions with incomplete cytokinesis, thereby generating a group of 16 interconnected spermatogonia. These further differentiate into spermatocytes that undergo meiosis to form spermatids and mature sperm [22,23]. Each GSC is surrounded by a pair of CySCs, which self-renew and give rise to cyst cells (CCs). The somatic CCs enclose GBs and provide signals necessary for germ cell maturation [24]. Specific gene expression patterns and markers can be used to identify both stem cell pools and differentiating daughters along the length of the testis [9].

In the absence of available antibodies specifically raised against *Drosophila* Tor, we used an existing fly line expressing a *Tor::GFP* fusion protein. The transgene consists of a large genomic clone containing *Tor* coding and regulatory sequences with a GFP tag inserted at the C-terminus, thereby providing a reporter designed to reflect endogenous protein expression [25]. RNAi-mediated knock down of Tor in germ cells efficiently reduced GFP signal in these cells, confirming that the GFP fluorescence indeed reflects the expression of GFP-tagged Tor (**Supp. Fig 1A-B**). Surprisingly, we observed that Tor::GFP is not uniformly expressed in young adult testes (**Fig 1A**). Indeed, cells at the tip of the testis, including GSCs (**Fig 1B**, yellow circles, arrowhead) and early spermatogonia (**Fig 1B**, magenta circles, arrow), exhibit higher levels of Tor::GFP, relative to mature spermatogonia (**Fig 1B**), while spermatocytes, identified as germ cells expressing high levels of Lamin C [26], have very low to undetectable Tor::GFP expression (**Fig 1C**). Interestingly, Tor::GFP expression varies in a similar pattern in the ovary, as female germ cells progress through differentiation during the first stages of oogenesis (**Supp. Fig 1C**), with higher expression detected in regions 1 and 2a of the germarium [27]. In addition, Tor::GFP progressively becomes restricted to the nucleus in cells expressing *bag of marbles* (*bam*) (**Fig 1B**). Taken together, these data suggest that Tor protein expression and localization varies with germ cell differentiation in males and females.

**Figure 1:**
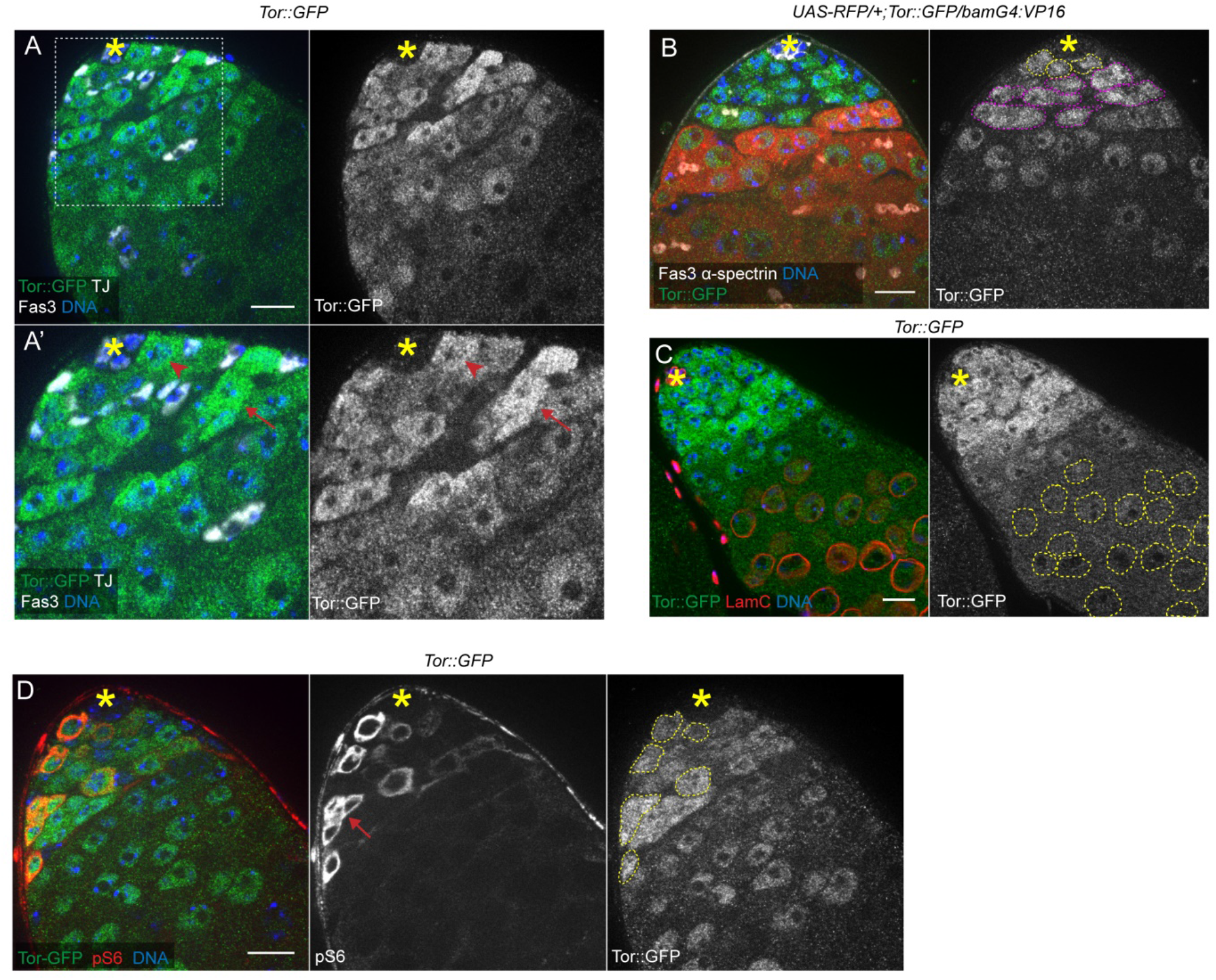
Tor expression and activity are higher in germline stem and progenitor cells. (A-A’) Expression pattern of Tor::GFP in an adult testis tip. Traffic-Jam (TJ) marks the nuclei of early cyst cells and Fasciclin 3 (Fas3) stains the membrane of hub cells. Germ cells are identified by the absence of TJ staining. A close-up of the region framed in (A) is shown in (A’). The red arrowhead points to an example of GSC and the red arrow indicates a spermatogonial cyst. (B) Tor::GFP expression pattern in a testis tip expressing RFP under the control of *bam-GAL4*. α-spectrin marks the fusome in germ cells and Fas3 is used to identify hub cells. GSCs, in contact with hub cells, are circled in yellow and early spermatogonia that do not express RFP are circled in magenta. (C) Tor::GFP expression pattern in a testis tip co-stained with LaminC (LamC). LamC is expressed at high levels in hub cells (asterisk) and in spermatocytes, circled in yellow. (D) Anti-phospho-dRpS6 (pS6) staining in the tip of a *Tor::GFP* testis. The red arrow points to an example of spermatogonial cyst positive for pS6. pS6-positive germ cells are circled in yellow dotted lines in the right panel showing Tor::GFP expression alone. In all panels, Tor::GFP is visualized by immunostaining with an anti-GFP antibody and asterisks indicate the location of the hub at the apical tip of the testis. Scale bars: 15µm.

In order to determine whether TOR is active in cells that express Tor::GFP, we took advantage of antibodies that specifically recognize phosphorylated *Drosophila* ribosomal protein S6 (dRpS6). Immunostaining with these antibodies was demonstrated previously to provide an accurate readout of TOR activity *in situ* in larval imaginal discs [28,29]. In adult testes, phosphorylated dRpS6 (hereafter referred to as pS6) was detected in a subset of GSCs and early germ cells (**Fig 1D**). As expected, pS6 was exclusively detected in germ cells expressing Tor::GFP and was absent in cells devoid of Tor::GFP, such as hub cells (**Fig 1D**, right panel). We validated pS6 as a reporter of TOR activity in the germline by feeding flies rapamycin, a potent mTORC1 inhibitor [30]. Upon rapamycin feeding, pS6 was largely lost in testes, indicating that pS6 is dependent on mTORC1 activity; Tor::GFP expression was not affected by feeding flies rapamycin (**Supp. Fig 1D-E**). In addition, pS6 levels appear to reflect genetically induced changes in mTORC1 activity in germ cells. Depletion of Iml1, a component of the GATOR1 complex that represses mTORC1 activity [31,32], induced a marked increase in the number of pS6-positive germ cells, when compared to depletion of *white* as a control (**Supp. Fig 1F-G**). By contrast, another validated TOR target, phosphorylated 4E-BP (p4E-BP) was not detected in germ cells by immunostaining (**Supp. Fig 1H**), even upon genetically induced upregulation of TOR activity (**Supp. Fig 1I**). However, overexpression of 4E-BP in germ cells led to a pattern of p4E-BP similar to pS6, suggesting that this may be due to low endogenous expression of 4E-BP (**Supp. Fig 1J**). Altogether, our data indicate that TOR is predominantly active in GSCs and early germ cells and that TOR activity is regulated at least in part through differential expression of the Tor kinase.

### Tor is required in adult GSCs and early germ cells for germline differentiation

In the adult testis, mTORC1 appears active in subsets of early germ cells (**Fig 1D**). In order to assess the role of mTORC1 activity in supporting germline homeostasis, we used RNA interference (RNAi) combined with the bipartite UAS/GAL4 system [33] to specifically deplete Tor kinase in GCSs and early germ cells. The *nanos-GAL4* ‘driver’ promotes expression in GSCs and early germ cells [34]; when combined with a ubiquitously expressed thermo-sensitive *GAL80*^*TS*^ allele [35] (hereafter referred to as *nanos*^*TS*^), this system permits depletion specifically in adult GSCs and early germ cells upon a shift to 29°C (cf. Methods).

Induction of *Tor*^*RNAi*^ under the control of *nanos*^*TS*^ led to a significant decrease in Tor::GFP expression (**Supp. Fig 1C**) and a decrease in pS6 staining specifically in Vasa^+^ germ cells (**Supp. Fig 2A-B**), confirming efficient depletion of Tor and subsequent loss of TOR activity in these cells. Conditional depletion of Tor kinase in adult male GSCs and early germ cells resulted in thinner testes (**Fig 2A-B**). However, germ cells were still present at the tip of the testis in contact with the hub (**Fig 2C-D**). In order to determine whether Tor-depleted germ cells were able to differentiate, we assessed the distribution pattern of α-spectrin to highlight the fusome, a membranous structure that is spherical in GSCs and GBs and forms a branched structure as spermatogonia undergo TA divisions [36,37]. Depletion of Tor in adult GSCs/early germ cells caused an accumulation of germ cells with a spherical fusome (**Fig 2C-D**), suggesting that decreased Tor may lead to an arrest in germ cell differentiation, prior to TA divisions. Alternatively, differentiating germ cells could de-differentiate, as has been described previously [38].

**Figure 2:**
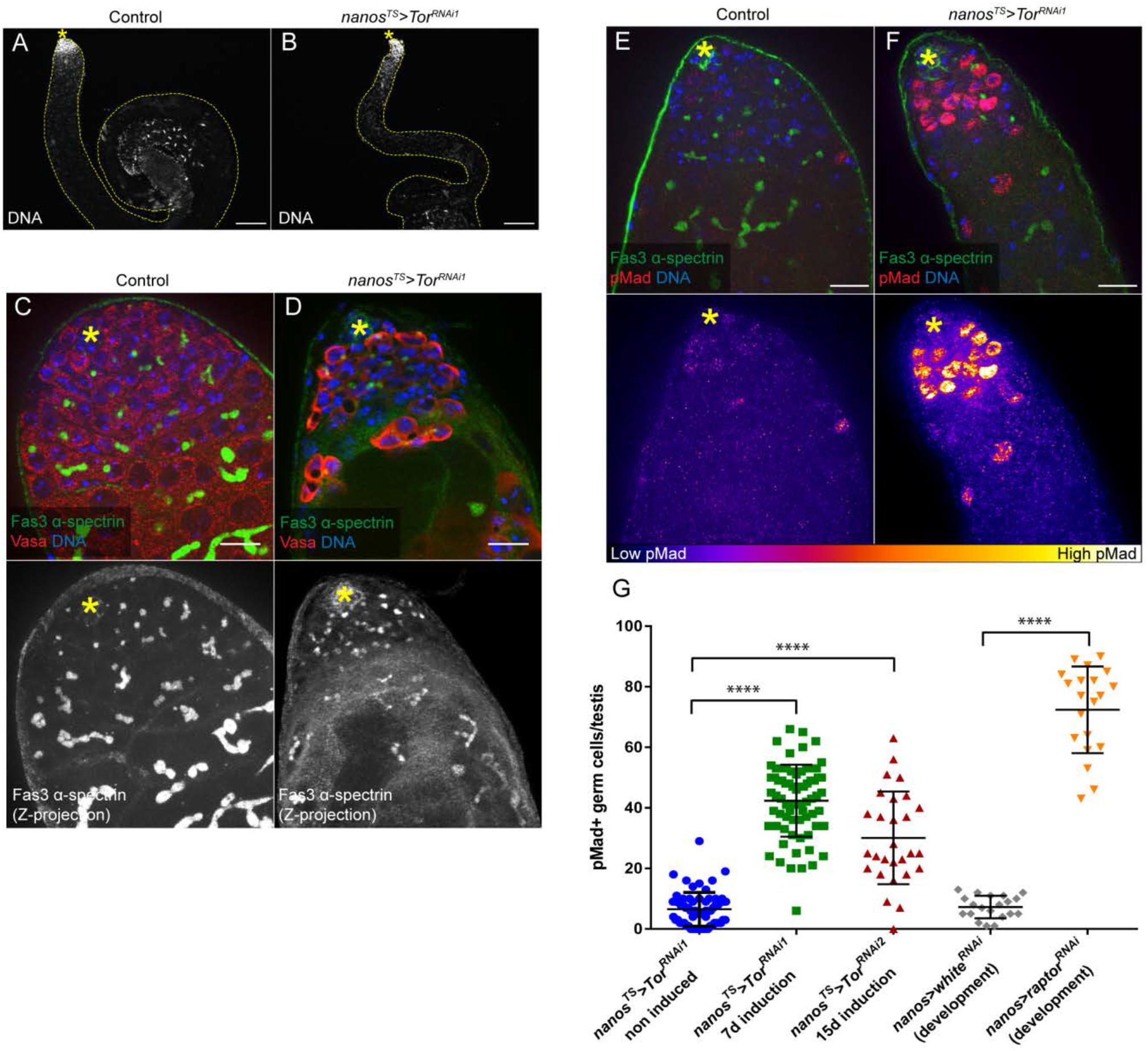
Downregulation of mTORC1 in early germ cells causes an accumulation of GSC-like germ cells. (A-B) Overall structure of single testes, visualized by staining nuclei with DAPI (DNA) and contoured with yellow lines in control (A) and upon early germ cell-specific depletion of Tor (B). Scale bar: 100µm. (C-D) Control (C) and Tor-depleted (D) testis tips with Vasa, α-spectrin and Fas3 stainings. Lower panels show maximum intensity projections of several images spanning the hub region (Z-projections). Scale bar: 15µm. (E-F) Control (E) and Tor-depleted (F) testis tips stained for phospho-Mad (pMad), α-spectrin and Fas3. Lower panels show pMad-associated fluorescence intensity represented as a color gradient with the lowest intensity in black and the brightest intensity in white. Arrowheads indicate pMad^+^ cyst cells. Scale bar: 15µm. (G) Number of pMad-positive germ cells per testis tip of the indicated conditions: nanos^TS^>Tor^RNAi1^ - non induced (n=66), nanos^TS^>Tor^RNAi1^ - 7 days induction (n=71), nanos^TS^>Tor^RNAi2^ - 15 days induction (n=28), nanos>white^RNAi^ - expressed throughout development (n=20) and nanos>raptor^RNAi^ - expressed throughout development (n=20). Means and standard deviations are shown. **** denotes statistical significance (p<0.0001) as determined with two-tailed unpaired t-tests. Testes shown in panels A to F are from flies of the genotype nanos-GAL4/+;UAS-Tor^TRiP-HMS00904^/tub-GAL80^ts^, either maintained at 18°C as a non-induced control (A, C and E) or raised at 18°C and shifted to 29°C at the adult stage for 7 days to induce Tor depletion in GSCs and early germ cells (B, D and F). In all panels, asterisks indicate the hub.

To further characterize the impact of Tor-depletion on germ cell differentiation, phosphorylation of Mothers against Dpp (Mad) was measured as a readout for Bone Morphogenetic Protein (BMP) pathway activity [39,40]. The BMP pathway is normally activated in GSCs by Dpp and Gbb ligands, secreted by somatic hub cells and CySCs, which regulates germline differentiation [41–43]. Depletion of Tor led to a dramatic increase in the number of germ cells positive for phosphorylated Mad (pMad) (**Fig 2E-G**). In addition, the intensity of pMad staining in Tor-depleted germ cells was markedly higher than in control germ cells (**Fig 2E-F**), and pMad positive germ cells were observed several cell diameters away from the hub. Taken together, these data support our observations that depletion of Tor results in accumulation of GSCs/GBs and suggest that mTORC1 negatively regulates BMP signaling in early germ cells.

Another RNAi transgene targeting a different region of *Tor* mRNA induced a similar block in germ cell differentiation and an accumulation of pMad positive germ cells, albeit upon a longer induction of RNAi expression (**Fig 2G; Supp. Fig 2C-D**). A similar phenotype was observed upon feeding flies with rapamycin (**Supp. Fig 2E-F**), as well as upon depletion of the mTORC1-specific subunit Raptor in GSCs and early germ cells throughout development (**Fig 2G; Supp. Fig 2F-G**).

Accumulation of GSC-like cells is not caused by stress or potential premature aging due to shifting the flies from 18°C to 29°C, as a control RNAi expressed under the control of *nanos*^*TS*^ at 29°C did not induce an increase in pMad^+^ germ cells, nor did aging of *nanos*^*TS*^*>Tor*^*RNAi*^ flies at 18°C for four weeks, which blocked induction of RNAi expression (**Supp. Fig 2I**). In addition, depletion of Tor in 4-16 cell spermatogonial cysts throughout development did not produce any obvious phenotype in the testis (**Supp. Fig 2J-K**), in agreement with the observation that Tor::GFP expression starts to decrease in cells that typically express *bag of marbles* (*bam*) (**Fig 1B**). Altogether, our results indicate that downregulation of mTORC1 activity in adult male GSCs and early germ cells causes a block at the earliest steps of germ cell differentiation, leading to the accumulation of germ cells harboring GSC-like features, including spherical fusomes and high BMP signaling.

### Tor depletion in GSCs and early germ cells leads to the accumulation of dysfunctional autolysosomes

Staining testes with antibodies against the germ cell marker Vasa revealed the presence of large areas devoid of Vasa expression in the cytoplasm of Tor-depleted germ cells (**Fig 3A-B**, arrowheads). Similar observations were made using all RNAi lines targeting Tor, as well as those targeting Raptor (**Supp. Fig 3A-B**), indicating that this phenotype correlates with mTORC1 downregulation in germ cells. By contrast, Vasa staining was uniform in the cytoplasm of spermatogonial cysts when Tor was depleted in 4-16 cell cysts (**Supp. Fig 2K**).

**Figure 3:**
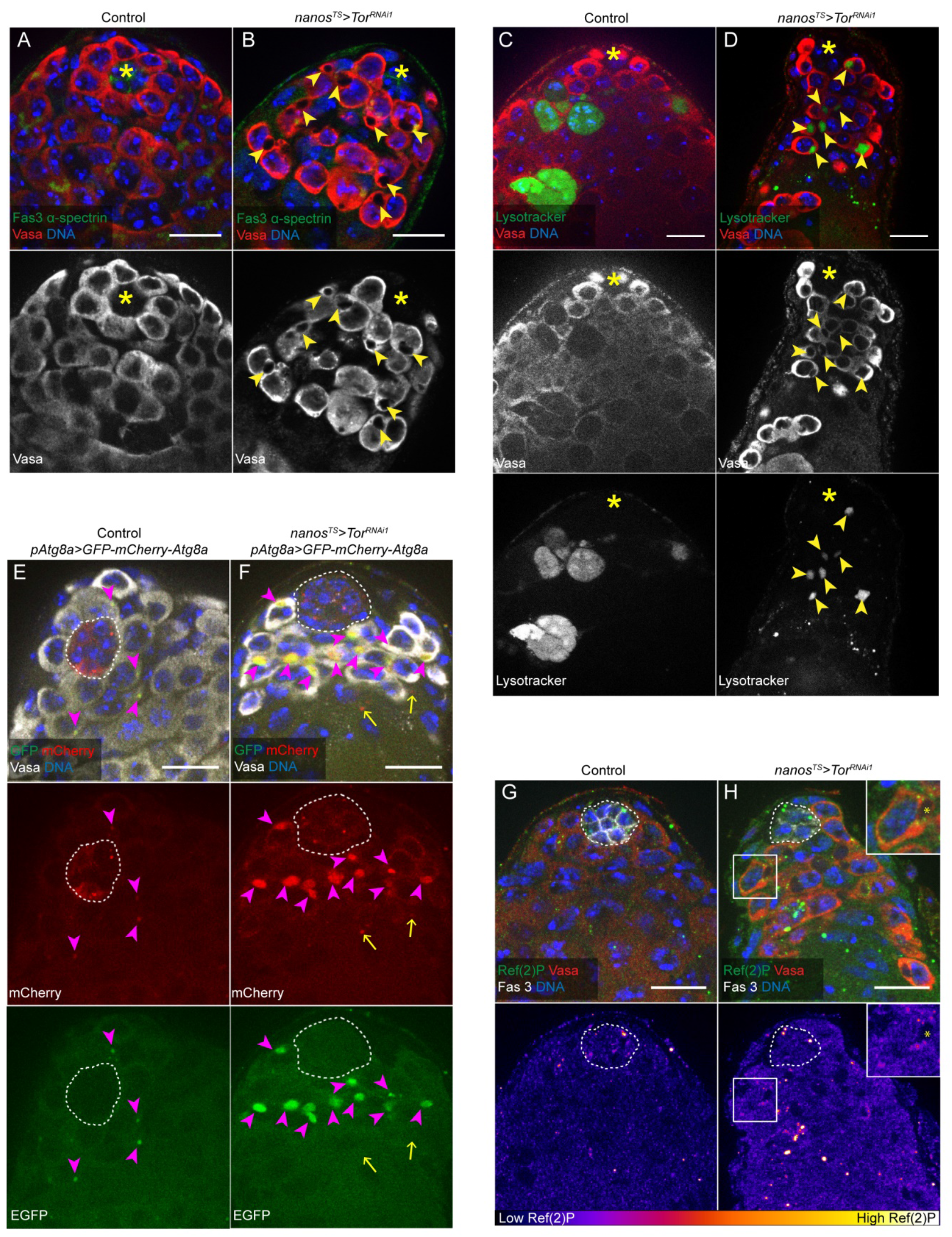
Tor depletion in GSCs/early germ cells leads to the accumulation of dysfunctional autolysosomes. (A-B) Control (A) and Tor-depleted (B) testis tips with Vasa, α-spectrin and Fas3 stainings. Arrowheads signal large cytoplasmic areas devoid of Vasa staining in Tor-depleted germ cells. (C-D) Control (C) and Tor-depleted (D) testis tips with Lysotracker Red and Vasa stainings. Arrowheads signal Vasa-negative and Lysotracker-positive cytoplasmic areas in Tor-depleted germ cells. (E-F) Control (E) and Tor-depleted (F) testis tips ubiquitously expressing the GFP-mCherry-Atg8a autophagic flux sensor. Magenta arrowheads signal yellow (GFP and mCherry-positive) Atg8a puncta in germ cells and the yellow arrows point to red (mCherry-positive and GFP-negative) Atg8a puncta in cyst cells (G-H) Control (G) and Tor-depleted (H) testis tips stained with Ref(2)P, Vasa and Fas3. The inset in H is a magnification of the framed germ cell, showing Ref(2)P aggregates in the cytoplasm, outside of the Vasa-devoid cytoplasmic are denoted with an asterisk. Lower panels show Ref(2)P-associated fluorescence intensity represented as a color gradient with the lowest intensity in black and the brightest intensity in white. Note that testes shown in panels (A-D) and (G-H) are from flies of the genotype *nanos-GAL4/+;UAS-Tor*^*TRiP-HMS00904*^*/tub-GAL80*^*ts*^ and testes in panels (E-F) are from flies of the genotype *nanos-GAL4/pAtg8a-GFP-mCherry-Atg8a;UAS-Tor*^*TRiP-HMS00904*^*/tub-GAL80*^*ts*^. Control corresponds to non-induced conditions (flies maintained at 18°C) and *nanos*^*TS*^*>Tor*^*RNAi1*^ corresponds to induced conditions (flies raised at 18°C and shifted to 29°C for 7 days at the adult stage). The hub is indicated with an asterisk in (A-D) and is delimitated by white dotted lines in (E-H). For all images, scale bar: 15µm.

Given the multiple roles of mTORC1 in repressing autophagy and lysosomal function (reviewed in [44,45]), we assessed whether these areas devoid of Vasa expression correspond to autophagic structures. Testes in which Tor was depleted from GSCs and early germ cells were stained with Lysotracker Red, a membrane-permeable vital dye that labels acidic compartments and is used to identify lysosomes [46]. In control testes, Lysotracker stained groups of spermatogonial cells, and the dye was distributed diffusely throughout the germ cells (**Fig 3C**), a pattern previously characterized in adult testes and corresponding to naturally occurring spermatogonial cell death [47,48]. By contrast, in Tor-depleted germ cells, Lysotracker consistently labeled cytoplasmic regions devoid of Vasa staining (**Fig 3D**, arrowheads), suggesting that these may be lysosomes. We did not observe diffuse Lysotracker staining as is observed in dying spermatogonia, supporting our hypothesis that the accumulation of early germ cells is the result of a defect in differentiation, rather than cell death in spermatogonia.

To determine whether these Lysotracker positive structures are autolysosomes, in which autophagic cargos are degraded, we assessed the expression of an mCherry-tagged *Atg8a* transgene, placed under the control of the *Atg8a* promoter [49], which permits visualization of all autophagic structures [46]. In control testes, mCherry-Atg8a formed small puncta that were primarily detected in hub cells and occasionally in CCs and spermatogonia, as previously reported [50] (**Supp. Fig 3C**). By contrast, upon depletion of Tor, mCherry-Atg8a formed large structures that filled the cytoplasmic regions devoid of Vasa staining (**Supp. Fig 3F**). Staining positive for both Lysotracker and Atg8a is indicative of autolysosomes. Moreover, as mCherry-Atg8a is expressed under the control of A*tg8a* promoter, our observations indicate that Tor depletion induces an upregulation of Atg8a expression in germ cells, consistent with recent reports showing that mTORC1 negatively regulates the expression of autophagy related genes, including *Atg8a*, by affecting mRNA processing [51,52].

The autolysosomes that form in germ cells upon Tor depletion appeared unusually large, which could reflect a defect in autophagic flux and lysosomal degradation. To test this hypothesis, we monitored the expression of a dual-tagged *GFP-mCherry-Atg8a* transgene, expressed under the control of *Atg8a* regulatory sequences [53]. GFP-mCherry-Atg8a appears yellow in autophagosomes; upon fusion with lysosomes, GFP fluorescence is more rapidly quenched by the acidic environment in the autolysosome, such that tagged Atg8a appears red. Therefore, we predicted that we would observe red and yellow puncta in control testes, but if autophagic flux is interrupted, the puncta will remain yellow. In control testes, GFP-negative and mCherry-positive Atg8a puncta were primarily observed in hub cells (**Fig 3E**, hub delimited by dashed line), indicating that autophagasomes present in these somatic cells are able to acidify, in agreement with previous findings [54]. The rare Atg8a puncta observed in germ cells in control testes appeared yellow (**Fig 3E**, arrowheads), suggesting that either these puncta correspond to autophagosomes that have not fused with lysosomes or that autolysosomes in wild-type germ cells do not properly acidify. In testes in which Tor was depleted in adult GSCs and early germ cells, red Atg8a puncta were observed in the hub (**Fig 3F**, circled) and in some CCs (**Fig 3F**, arrows). Remarkably, large yellow Atg8a vesicles accumulated in the cytoplasm of germ cells (**Fig 3F**, arrowheads). These data suggest that autolysosomes that form upon targeted depletion of Tor in adult early germ cells do not properly acidify. A ubiquitously expressed GFP-LAMP fusion protein [55,56], which is targeted to lysosomes where hydrolases normally degrade GFP, was also detected in these vesicles (**Supp. Fig 3E-F**), further supporting the finding that acidification either does not take place or is not efficient.

Ref(2)P, or p62 in mammals, is an intracellular receptor that binds ubiquitinated proteins and targets them for selective autophagic degradation [57]. As such, accumulation of Ref(2)P is commonly used as a readout for defects in autophagy [46]. Upon depletion of Tor in GSCs and early germ cells, we observed an increase in Ref(2)P aggregates in germ cells (**Fig 3G-H**). This is in agreement with a defect in autophagic flux leading to the accumulation of un-degraded proteins marked with Ref(2)P. Therefore, consistent with a known role for mTORC1 for repressing autophagy, our observations indicate that Tor depletion leads to an increase in autophagic structures in GSCs and early germ cells.

### Down-regulation of Tor in adult GSCs and early germ cells induces ectopic TOR activity in the soma and premature differentiation of cyst cells

When verifying that pS6 was efficiently lost in adult GSCs/early germ cells upon Tor depletion, a concomitant increase in pS6 signal was observed in the adjacent somatic CC compartment (**Supp. Fig 2B**). Indeed, the number of pS6^+^ early CCs, identified by the expression of Traffic Jam (TJ), was significantly higher when Tor was depleted in adult GSCs/germ cells (**Fig 4A-C**). This increase in pS6 reflects *bona fide* mTORC1 activity, as it is lost upon rapamycin treatment (**Supp. Fig 4A-B**). We also observed a similar increase in TOR activity when *raptor* was depleted in early germ cells, throughout development (**Supp. Fig 4C-D**). Therefore, downregulation of mTORC1 activity in the germline induces non autonomous activation of mTORC1 in the neighboring CCs.

**Figure 4:**
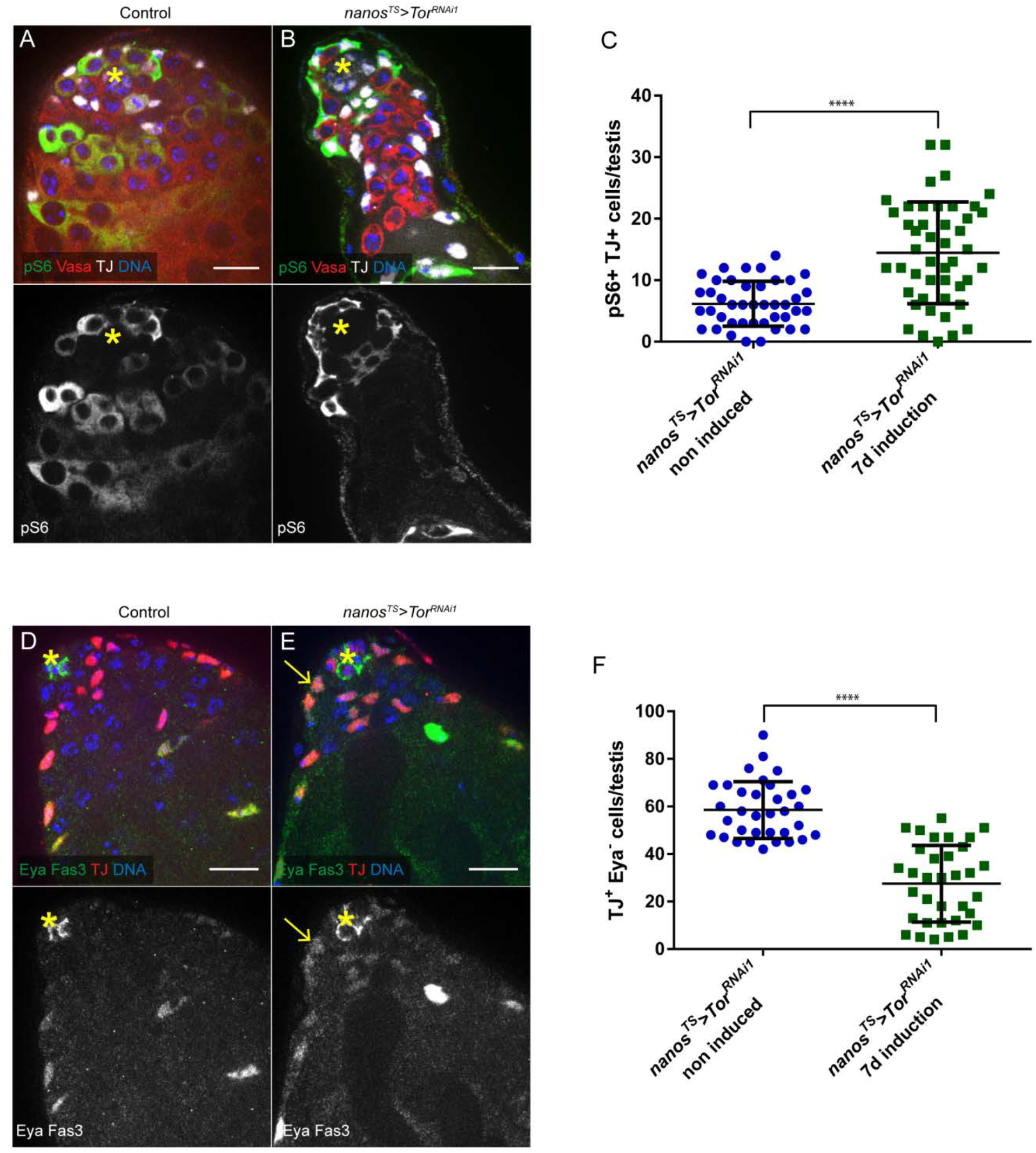
Down-regulation of Tor in adult GSCs/early germ cells induces ectopic TOR activity in the soma and premature differentiation of cyst cells. (A-C) Down-regulation of Tor in germ cells induces a concomitant increase in TOR activity in neighboring cyst cells. (A-B) Control (A) and Tor-depleted (B) testis tips with pS6, Vasa and TJ stainings. (C) Number of pS6- and TJ-double positive cells per testis tip in the indicated conditions: *nanos*^*TS*^*>Tor*^*RNAi1*^ - non induced (n=42) and *nanos*^*TS*^*>Tor*^*RNAi1*^ - 7days induction (n=46). Means and standard deviations are shown. **** denotes statistical significance (p<0.0001) as determined with two-tailed unpaired t-tests. (D-F) Down-regulation of Tor in germ cells leads to a loss of early cyst cells. (D-E) Control (D) and Tor-depleted (E) testis tips with Eya, TJ and Fas3 stainings. The arrow in (E) points to an example of Eya-positive cyst cell in proximity to the hub. Number of early cyst cells (TJ positive and Eya negative cells) per testis tip of the indicated conditions: *nanos*^*TS*^*>Tor*^*RNAi1*^ - non induced (n=34) and *nanos*^*TS*^*>Tor*^*RNAi1*^ - 7days induction (n=34). Means and standard deviations are shown. **** denotes statistical significance (p<0.0001) as determined with two-tailed unpaired t-tests. Testes shown in panels (A-B) and (D-E) are from flies of the genotype *nanos-GAL4/+;UAS-Tor*^*TRiP-HMS00904*^*/tub-GAL80*^*ts*^. Control corresponds to non-induced conditions (flies maintained at 18°C) and *nanos*^*TS*^*>Tor*^*RNAi1*^ corresponds to induced conditions (flies raised at 18°C and shifted to 29°C for 7 days at the adult stage). The hub is indicated with an asterisk. For all images, scale bar: 15µm.

mTORC1 activity is required for differentiation of CCs, and CC-specific downregulation of mTORC1 leads to an accumulation of early CCs [21,58]. Thus, we monitored the differentiation of the CC lineage in testes in which Tor is depleted in adult early germ cells. In the somatic lineage, the transcription factor TJ is expressed in CySCs and CCs that surround spermatogonia undergoing TA divisions [59], while late-stage CCs in contact with differentiating spermatocytes express high levels of the transcription factor Eyes absent (Eya) [60]. Therefore, the number of early CCs can be quantified by counting the number of cells expressing TJ but expressing low or no Eya [50]. Upon depletion of Tor in GSCs and early germ cells, a marked decrease in the number of TJ+ Eyaearly CCs was observed (**Fig 4F**). In addition, Eya+ cells were observed closer to the testis tip (**Fig 4D-E**), including in contact with the hub (**Fig 4E**, arrow). The loss of early CCs, together with the observation that more mature CCs are located at the testis tip, suggest that CCs may differentiate prematurely or at a higher rate upon downregulation of TOR in germ cells and the subsequent non-autonomous upregulation of TOR in the soma. Of note, the Hedgehog target gene Patched (Ptc), which is also used as a marker for CySCs [61], was expressed in somatic cells directly in contact with the hub (**Supp Fig 4E-F**, arrowheads), suggesting that Hedgehog signaling is not affected. Strikingly, mature CCs are found adjacent to Tor-depleted germ cells that exhibit GSC-like features (**Fig 2**), indicating that somatic and germline differentiation appear to be uncoupled. Rapamycin treatment led to a complete loss of Eya expression (**Supp Fig 4G-H**), both in control testes, as previously reported [21,58], and in testes expressing *Tor*^*RNAi*^ in GSCs and early germ cells (**Supp Fig 4I-J**). Hence, the premature expression of Eya observed in this condition is dependent on mTORC1 activity.

## Discussion

Our findings highlight an essential role for the TOR pathway in regulating adult GSC behavior in the *Drosophila* testis. Germ cell specific RNAi-mediated depletion of the Tor kinase prevented germ cell differentiation and induced an accumulation of germ cells expressing markers of GSCs. In addition, cells depleted for Tor exhibited abnormally elevated BMP signaling, upregulated autophagy and an accumulation of dysfunctional autolysosomes. Depletion of Tor in GSCs and early germ cells also led to non-autonomous activation of mTORC1 in neighboring CCs and premature differentiation of the CC lineage. Tor has previously been shown to play a role in regulating proliferation and differentiation in a variety of stem cell systems, including the fly female germline [17], and our data indicate that Tor plays a similar role in early germ cells of the fly testis. Using transgenic flies expressing Tor::GFP under the control of *Tor* regulatory sequences, we observed that the Tor kinase is differentially expressed among early germ cells in the testis, suggesting that not all early germ cells may be as permissive to upstream signals modulating TOR activity. Consistent with the variation in expression, we found that mTORC1 activity was restricted to Tor::GFP expressing germ cells, using pS6 as readout. Of note, p4E-BP, which is commonly used as a reporter for Tor activity, was not readily detected in germ cells, although overexpression of 4E-BP in germ cells did result in p4E-BP staining (**Supp Fig1J**), suggesting that 4E-BP may not be present at adequate levels in male germ cells under homeostatic conditions to be a primary target. Both pS6 and p4E-BP were detected in early CCs [21], consistent with the well-characterized role for Tor in the somatic lineage in the testis (**Figs 4A, 4C**, and **Supp Fig 1H**). Therefore, our observations highlight the importance of assessing the phosphorylation of different targets as readouts of TOR activity.

Similar to depletion of Tor in GSCs and early germ cells, germline-specific depletion of Raptor and ubiquitous inhibition of mTORC1 through rapamycin feeding led to an accumulation of germ cells harboring GSC-like features, including spherical fusomes and high levels of nuclear pMad, indicating signaling via the BMP pathway (**Fig 2**). Interestingly, mTORC1 was shown to be a negative regulator of BMP signaling in other models investigating progenitor cell differentiation [62–65]. For example, in the female *Drosophila* germline, downregulation of BMP signaling upon hyperactivation of TOR in GSCs contributes to premature differentiation [18]. In mammals, mTORC1 downregulates BMP signaling to promote hair follicle stem cell activation [66] and to promote the transition from early-stage oligodendrocyte precursor cells to immature oligodendrocytes in the central nervous system [65]. Thus, down-regulation of BMP signaling appears to be a conserved mechanism through which mTORC1 regulates the transition into a differentiation program. The mechanisms involved in this process remain to be elucidated. Although we find that BMP signaling is upregulated upon mTORC1 inhibition in GSCs, hyperactivation of BMP signaling in GSCs is not sufficient to prevent the transition to TA divisions [42,43,67]. Therefore, this is not likely to be the primary mechanism underlying the block in differentiation in male germ cells.

Depletion of Tor led to an accumulation of dysfunctional autolysosomes, together with Ref(2)P^+^ aggregates, suggesting that TOR may play an important role in regulating proteostasis in GSCs and early germ cells via repression of the autophagy pathway. Interestingly, BMP signaling has been shown previously to be a positive regulator of autophagy [68,69]. Therefore, it is possible that, in addition to the direct effects of Tor depletion on autophagy, upregulation of the BMP pathway in Tor-depleted germ cells also induces autophagy, leading to unusually large autolysosomes that appear to be stalled. Our laboratory recently reported that RNAi-mediated depletion of Atg proteins in GSCs and early germ cells does not have a significant impact on GSC maintenance or germline differentiation [50]. Therefore, we hypothesize that autophagy may be actively repressed by TOR signaling in GSCs and early germ cells under homeostatic conditions. We also find that downregulation of mTORC1 in GSCs and early germ cells led to increased mTORC1 activity in neighboring CCs, which correlated with a loss of early CCs through premature differentiation. This is reminiscent of studies in mice reporting that caloric restriction (CR) leads to inhibition of mTORC1 in the Paneth support cells of intestinal crypts and a concomitant increase in mTORC1 activity in adjacent intestinal stem cells, promoting their proliferation [70,71]. Of note, in the intestine, CR induces an increase in pS6 but does not affect p4E-BP, further highlighting the importance of assessing multiple readouts for TOR activity. The mechanisms underlying this cross-talk in the testis remain to be elucidated. One possibility is that metabolites released from germ cells, such as degradation products resulting from autophagy activation upon mTORC1 inhibition, act as signals to activate mTORC1 in neighboring cells. Indeed, autophagy can act as an upstream regulator of TOR [45], and it was previously shown that upon extended inhibition of mTORC1, the recycling of cellular macromolecules through autophagy can stimulate a feedback mechanism leading to mTORC1 reactivation [72]. A similar pathway was also described in the context of paligenosis, a process by which mature cells de-differentiate and acquire stem cell features in response to injury. Following injury, mTORC1 is first inhibited, leading to upregulation of the autophagic and lysosomal machineries, followed by subsequent reactivation of mTORC1 in a lysosome-dependent manner [73,74]. In our model, mTORC1 cannot be reactivated in GSCs and early germ cells as the Tor kinase is depleted in these cells. Therefore, degradation products generated by autophagy in Tor-depleted germ cells could be secreted via lysosomal exocytosis or other forms of non-conventional secretion that are dependent on autophagy [75] and signal to activate mTORC1 in neighboring CCs. In line with this hypothesis, a recent study showed that mTORC1 inhibition stimulates unconventional protein secretion [76].

In addition, the fact that germline differentiation is delayed or arrested, while somatic cell differentiation appears to be accelerated, indicates that there is uncoupling of the differentiation of these two lineages, which is reminiscent of the phenotypes induced by disruption of EGFR signaling. The EGF ligand Spitz is produced by GSCs to activate EGFR signaling in adjacent somatic cells [77,78], and under homeostatic conditions, EGFR activity is required autonomously for CySC maintenance. Similar to Tor depletion in GSCs and early germ cells, loss of function mutations in either *Egfr* [77] or the downstream effector *raf* [79] cause an accumulation of early germ cells, as well as somatic cells expressing differentiation markers, such as Eya, close to the hub [50,79]. Here, we find that depleting Tor in GSCs causes an activation of TOR in neighboring CCs, which appears to contribute to premature differentiation. In a previous study, we showed that EGFR and TOR signaling act antagonistically in CCs, as EGFR stimulates autophagy to control early CC behavior and TOR suppresses autophagy to allow CC differentiation [50]. It is not known whether EGFR contributes to TOR inhibition in early CCs. Based on our observations, one interesting hypothesis is that TOR could be required for normal production of Spitz by germ cells; if so, Tor depletion in GSCs would alter EGFR signaling in neighboring CCs. However, it may be that metabolites secreted from the Tor-depleted germ cells act together with altered EGFR signaling to disrupt tissue homeostasis in this model.

In summary, our analysis demonstrates that mTORC1 is required for the differentiation of male GSCs under homeostatic conditions, pointing to a conserved role for mTORC1 in regulating the transition of progenitor cells to a differentiation program in a number of adult stem cell systems [16]. In particular, studies in the mouse testis reported a similar correlation between mTORC1 activity and differentiation within the spermatogonial progenitor pool [80]. In the mouse testis, rapamycin treatment was also shown to block spermatogonial differentiation, leading to an accumulation of undifferentiated spermatogonia [80–82] that seem to harbor enhanced sensitivity to niche cytokines [80], while rapamycin analogs were associated with reversible male infertility in human [83,84]. Thus, mTORC1 requirement for spermatogenesis and germ cell differentiation appears to be conserved. Although it is not known if Tor is also differentially expressed in these cells, these findings raise the possibility that Tor expression may be developmentally controlled through conserved mechanisms.

## Acknowledgements

The authors thank Margaret Fuller (Stanford University), Thomas Neufeld (University of Minnesota), David Walker (University of California, Los Angeles), Helmut Krämer (University of Texas Southwestern), the Bloomington Drosophila Stock Center and the Vienna Drosophila Resource Center for fly stocks; Aurelio Teleman (German Cancer Research Center), Jongkyeong Chung (Seoul National University), Dorothea Godt (University of Toronto), David Walker (University of California, Los Angeles) and the Developmental Studies Hybridoma Bank for antibodies. We acknowledge members of the Jones laboratory for helpful discussions, Utpal Banerjee (University of California, Los Angeles) for providing access to laboratory space and reagents, the UCLA Eli and Edythe Broad Center of Regenerative Medicine and Stem Cell Research Training Program for support, and NIH: AG040288, AG052732, GM135767 (DLJ).

## Author Contributions

M.C. designed, performed, analyzed experiments and wrote the manuscript. C.D. designed, performed, and analyzed experiments. A.K. performed experiments. D.L.J. designed and analyzed experiments and wrote the manuscript.

## Materials and Methods

### Fly stocks and genetics

The *Drosophila* transgenes used in this study are listed in **Table 1**. All fly stocks were raised on standard cornmeal medium at 25°C, with the exception of crosses with the *nanos*^*TS*^ driver. Transgene expression in germ cells throughout development was achieved using *nanos-GAL4:VP16* or *bam-GAL4:VP16* at 25°C. Early adult specific depletion in early germ cells using the *nanos*^*TS*^ driver was achieved by raising flies at 18°C and shifting the adult progeny to 29°C, for either 7 days (*Tor*^*RNAi1*^) or 15 days (*Tor*^*RNAi2*^) to induce expression of the RNAi transgenes. We noted that this system had variable efficiency depending on the RNAi transgenes used. For example, expression of *raptor*^*RNAi*^ in adult germ cells using *nanos*^*TS*^ did not cause any reduction in pS6 staining (not shown), indicating that Raptor is likely not depleted at a sufficiently high level to cause a reduction in TOR activity, whereas its expression throughout development using *nanosGAL4:VP16* efficiently led to a loss of pS6 in germ cells (**Supp. Fig 4D**).

**Table 1.**
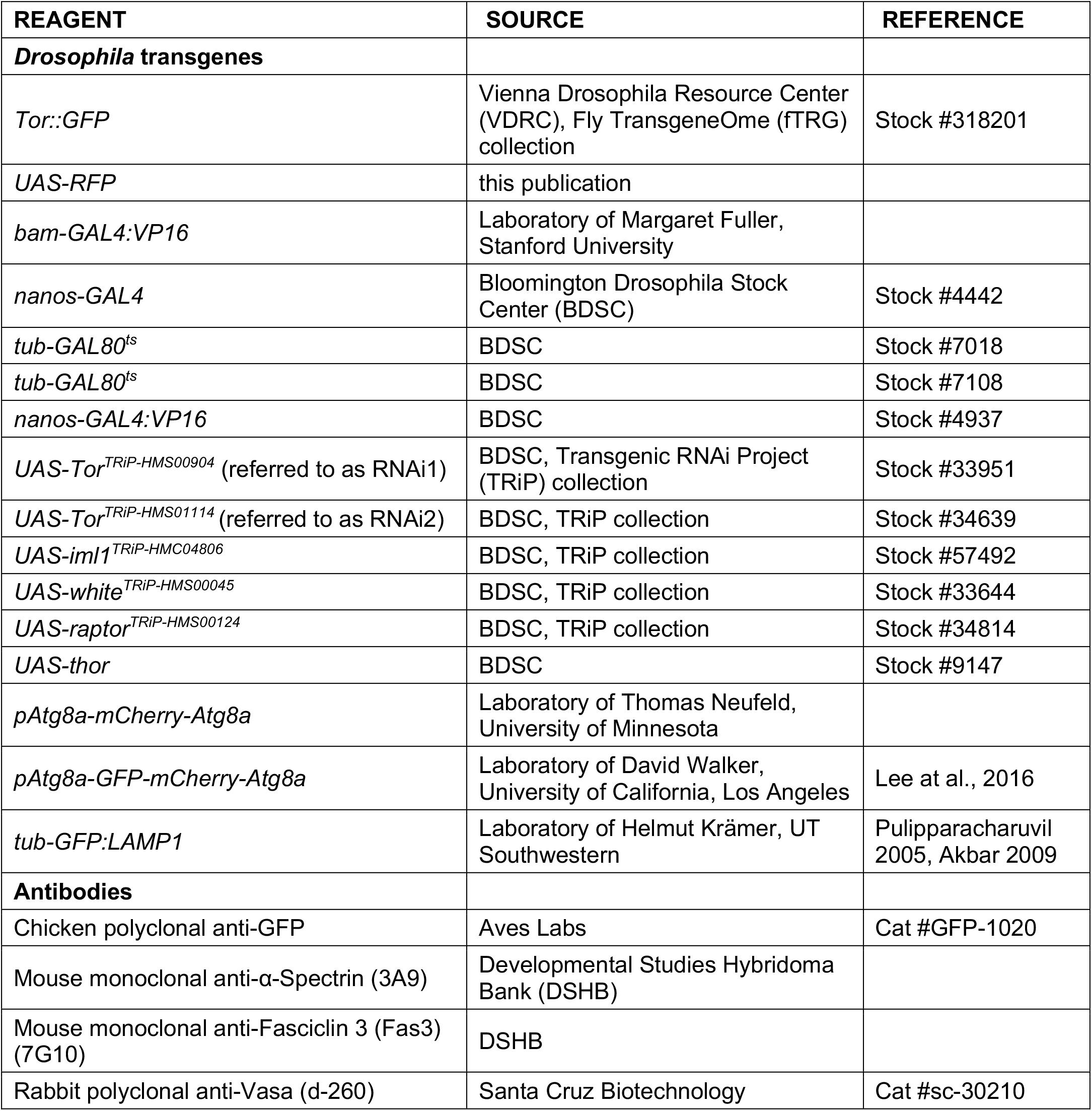
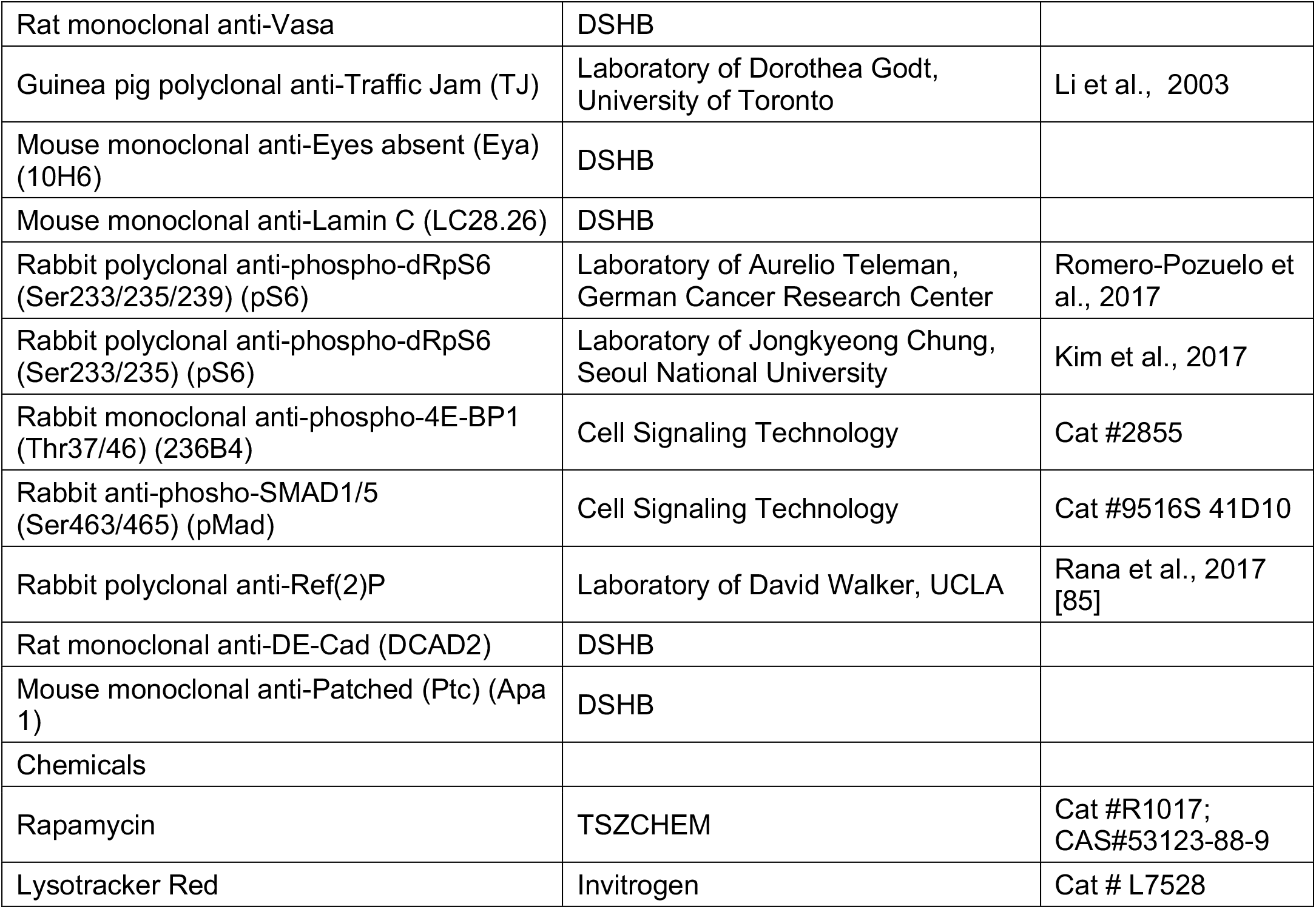
List of fly stocks and reagents.

### Rapamycin feeding

Rapamycin feeding was performed as previously described [21]. 100µl of a 4mM Rapamycin (TSZ Chem, Cat# R1017) stock solution in ethanol was added to the surface of standard diet food and let air dried. Similarly, control food was prepared by adding 100µl of ethanol to the surface of standard diet food. Flies were transferred to fresh rapamycin- or ethanol-treated food every 2-3 days, for the period of time indicated for each experiment.

### Immunostaining

Adult testes were dissected in PBS and fixed in 4% paraformaldehyde for 30 minutes, then permeabilized by two 15 minutes washes in 0.3% Sodium Deoxycholate and washed for 15 minutes in 0.1% PBT (1x PBS, 0.1% Triton X-100). Tissues were incubated in 3% Bovine Serum Albumine (BSA)-0.1% PBT blocking solution for at least 30 minutes. Primary antibodies (listed in **Table 1**) were diluted in blocking solution as follow: chicken anti-GFP (1:1000), mouse anti-α-Spectrin (1:20), mouse anti-Fas3 (1:100), rabbit anti-Vasa (1:100), rat anti-vasa (1:25), guinea pig anti-TJ (1:3000), mouse anti-Eya (1:20), mouse anti-LaminC (1:20), rabbit anti-phospho-dRpS6 Ser 233/235/239 (1:200), anti-phospho-dRpS6 Ser 233/235 (1:400), rabbit anti-phospho-4E-BP1 Thr37/46 (1:500), rabbit anti-phospho-SMAD1/5 Ser463/465 (1:100), rabbit anti-Ref(2)P (1:200), rat anti-DE-Cad (1:20) and mouse anti-Patched (1:500). Following overnight incubation at 4°C with primary antibodies, samples were washed with PBT three times for 15 minutes, incubated in Alexa-conjugated secondary antibodies (Invitrogen, 1:500 in blocking solution), washed with PBT three times for 15 minutes and mounted in Vectashield® (Vector laboratories) with DAPI.

For Lysotracker Red staining, testes were dissected in PBS and incubated for 30 minutes in Lysotracker Red solution (in 1:1000 in PBS) prior to fixation in 4% paraformaldehyde for 20 minutes and continuation with the immunostaining protocol as described above, maintaining the tissues in the dark.

### Microscopy and image analysis

Fluorescent images were acquired with a Carl Zeiss Axio Vert.A1 inverted light microscope using a 63x oil-immersion objective. Images were processed using ZEN digital imaging and analyzed with the ImageJ software. All images shown are representative images of at least three independent experiments.

Quantitative experiments were evaluated for statistical significance using the software Graphpad Prism v6.0, after verifying the normality of values and equivalence of variances. Graphics display means with standard deviations and the statistical differences between control and test samples were assessed using unpaired t-tests. Statistical significance is denoted as * for p < 0.05, ** for p < 0.01, *** for p < 0.001, **** for p < 0.0001 and n.s (not significant) for p > 0.05. The number of values in each condition is indicated in the figure legends.

---

**Supplementary Figure 1:**
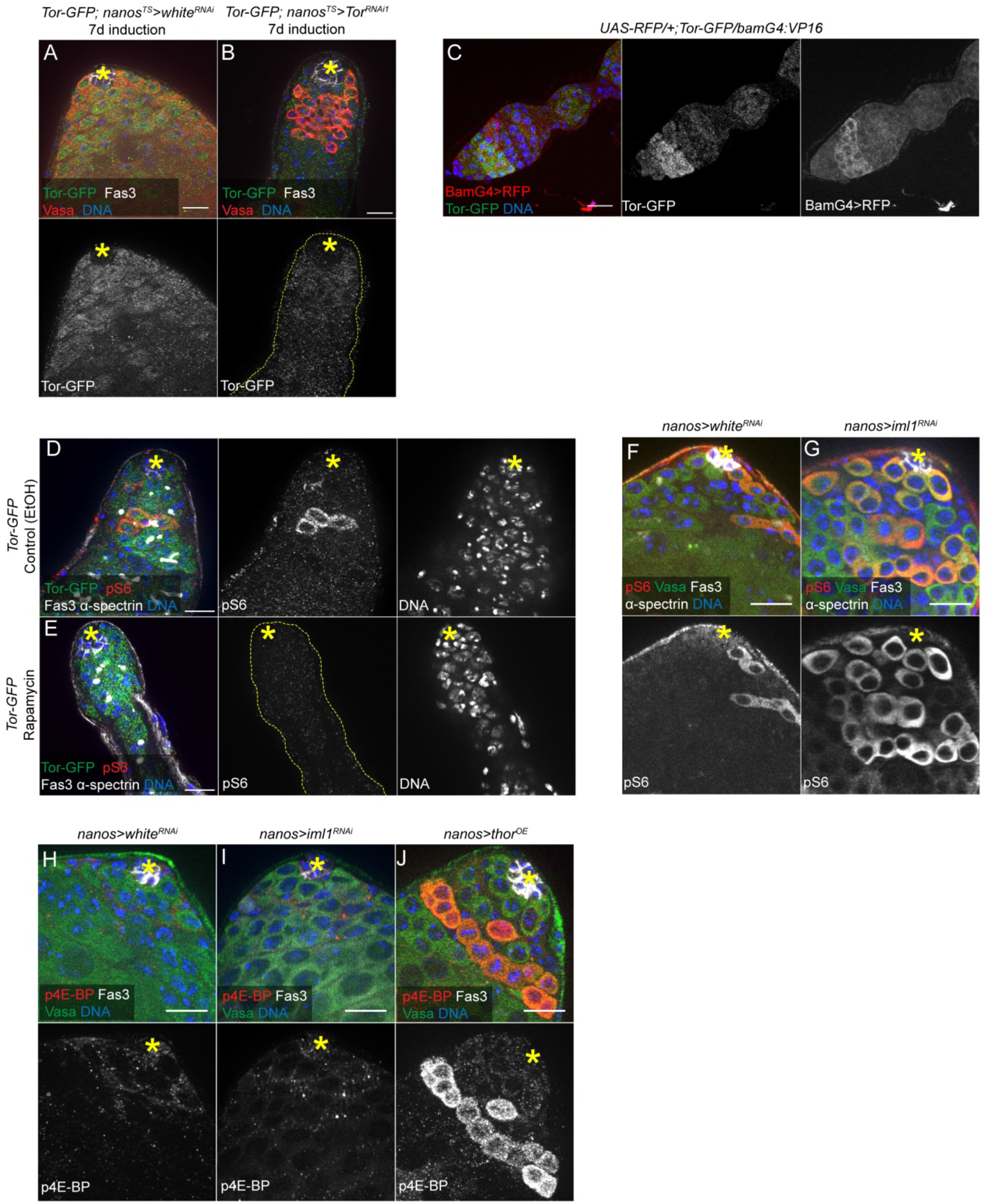
(A-B) Tor::GFP expression strongly decreases upon downregulation of Tor in GSCs and early germ cells. Images show examples of testis tips from control (A) (*nanos-GAL4/tub-GAL80*^*ts*^*;UAS-white*^*TRiP-HMS00045*^*/Tor::GFP*) and Tor-depleted (B) (*nanos-GAL4/tub-GAL80*^*ts*^*;UAS-Tor*^*TRiP-HMS00904*^*/Tor::GFP*) animals raised at 18°C and shifted to 29°C for 7 days at the adult stage to induce expression of the RNAi transgenes, with GFP, Fas3, Vasa and DNA stainings. In the lower (B) panel, the testis is contoured with dotted lines. (C) Tor::GFP expression pattern in the first stages of oogenesis. Image shows an example of germarium with one egg chamber from an ovary of a fly carrying the *Tor::GFP* transgene and expressing RFP under the control of *bam-GAL4*, stained with an anti-GFP antibody and DAPI. (D-E) Rapamycin treatment leads to a loss of pS6 staining without change in Tor::GFP expression. *Tor::GFP* flies were placed on food containing either ethanol as a control (D) or rapamycin (4mM in ethanol) (E) for 5 days. Images show testis tips with GFP, pS6, Fas3, α-spectrin and DNA stainings. Dotted lines delimitate the contour of the testis shown in (E). (F-G) Genetically induced hyperactivation of mTORC1 leads to an increase in pS6-positive cells. Images show examples of testis tips from control (F) (*nanos-GAL4:VP16/UAS-white*^*TRiP-HMS00045*^) animals, with pS6, Vasa, Fas3, α-spectrin and DNA stainings. (H-J) Phosphorylation of 4E-BP does not respond to variations in TOR activity in male germ cells. Images show testis tips from control (H) (*nanos-GAL4:VP16/UAS-white*^*TRiP-HMS00045*^), Iml1-depleted (I) (U*AS-iml1*^*TRiP-HMC04806*^*/+;nanos-GAL4:VP16/+*) and 4E-BP-overexpressing (J) (*UAS-thor/+;nanos-GAL4:VP16/+*), with p4E-BP, Fas3, Vasa and DNA stainings. Asterisks denote the hub. All scale bars: 15µm.

**Supplementary Figure 2:**
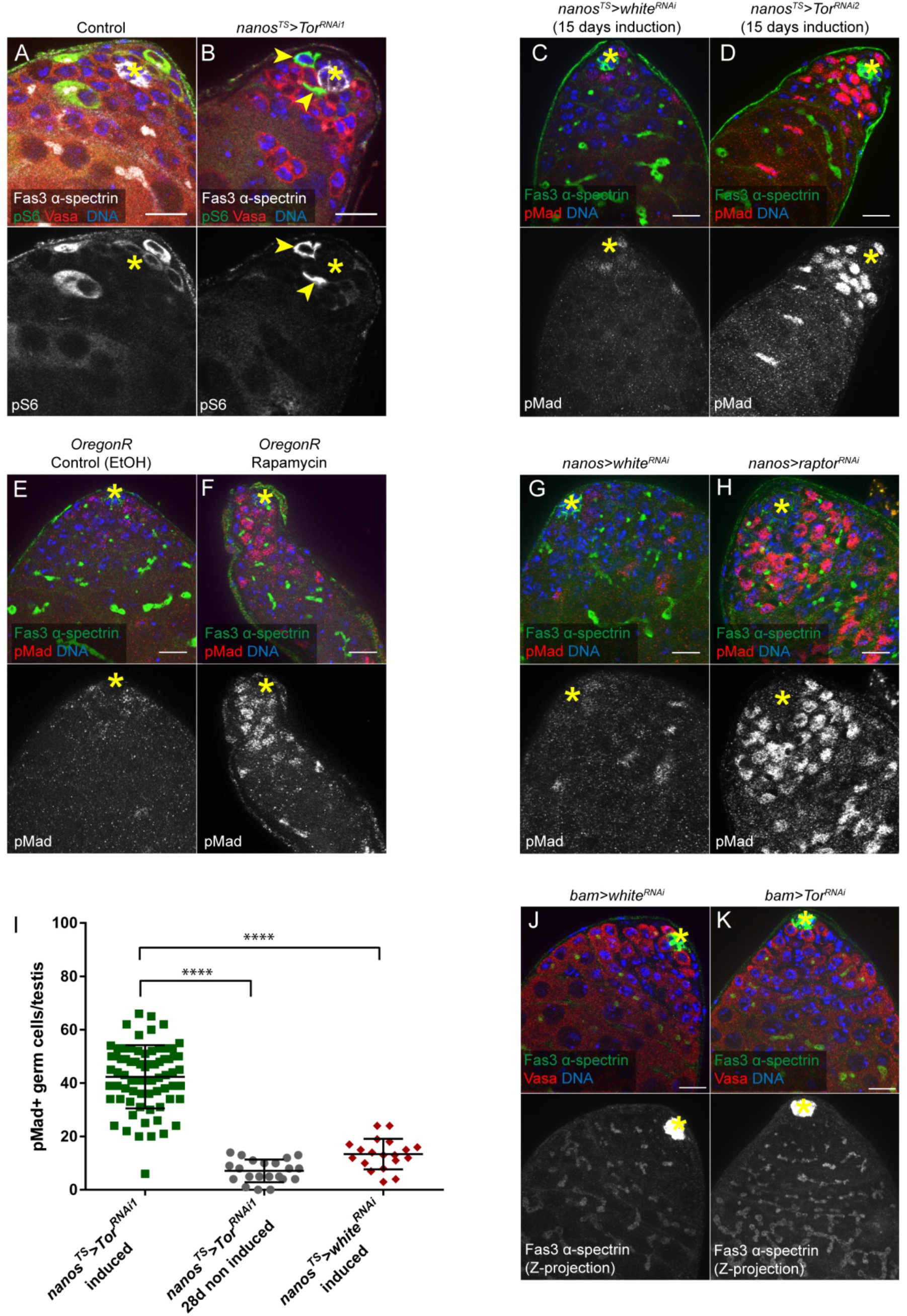
pS6. Images show testis tips from nanos-GAL4/+;UAS-Tor^TRiP-HMS00904^/tub-GAL80^ts^ animals raised at 18°C and either maintained at 18°C (A) (Control) or shifted to 29°C at the adult stage for 7 days prior to dissection (B) (nanos^TS^>Tor^RNAi1^), with pS6, Traffic Jam (TJ), Vasa and DNA stainings. pS6 positive cells are contoured in yellow in panels showing individual stainings. (C-D) RNAi-mediated depletion of Tor with RNAi2 induces the same phenotype as with RNAi1, with a longer induction time. Images show testis tips from nanos-GAL4/+;UAS-white^TRiP-HMS00045^ /tub-GAL80^ts^ (C) and nanos-GAL4/+;UAS-Tor^TRiP-HMS01114^/tub-GAL80^ts^ (D) animals raised at 18°C and shifted to 29°C at the adult stage for 15 days prior to dissection, with Fas3, α-spectrin, pMad and DNA stainings. (E-F) Rapamycin treatment.induces an increase in pMad-positive germ cells with a dotted spectrosome. Wild-type OregonR flies were placed on food containing either ethanol as a control (E) or rapamycin (4mM in ethanol) (F) for 7 days. Images show testis tips with Fas3, α-spectrin, pMad and DNA stainings. (G-H) RNAi-mediated depletion of Raptor throughout development causes an increase in pMad-positive cells with a dotted spectrosome. Images show testis tips from nanos-GAL4:VP16/UAS-white^TRiP-HMS00045^and nanos-GAL4:VP16/UAS-raptor^TRiP--HMS00124^ (H) animals, with Fas3, α-spectrin, pMad and DNA stainings. (I) Number of pMad-positive germ cells per testis tip of the indicated conditions: nanos^TS^>Tor^RNAi1^ - 7 days non induced (n=66), nanos^TS^>Tor^RNAi1^ - 7 days induction (n=71), nanos^TS^>Tor^RNAi2^ - 15 days induction (n=28), nanos^TS^>Tor^RNAi1^ - 28 days non induced (n=21) and nanos^TS^>white^RNAi^ - 10-15 days induction (n=19). Means and standard deviations are shown. **** denotes statistical significance (p<0.0001) as determined with two-tailed unpaired t-tests. Data from nanos^TS^>Tor^RNAi1^ - induced are the same as in Figure 2G and are ––shown here for comparison purpose. (J-K) Tor depletion in later stages of spermatogonial differentiation using bam-GAL4 does not have any obvious phenotype. Images show testis tips from bam-GAL4:VP16/UAS-white^TRiP-HMS00045^ (G) and bam-GAL4:VP16/UAS-raptor^TRiP--HMS00124^ (H) animals, with Fas3, α-spectrin, Vasa and DNA stainings. Lower panels show maximum intensity projections of several images spanning the hub region (Z-projections). Asterisks denote the hub. All scale bars: 15µm.

**Supplementary Figure 3:**
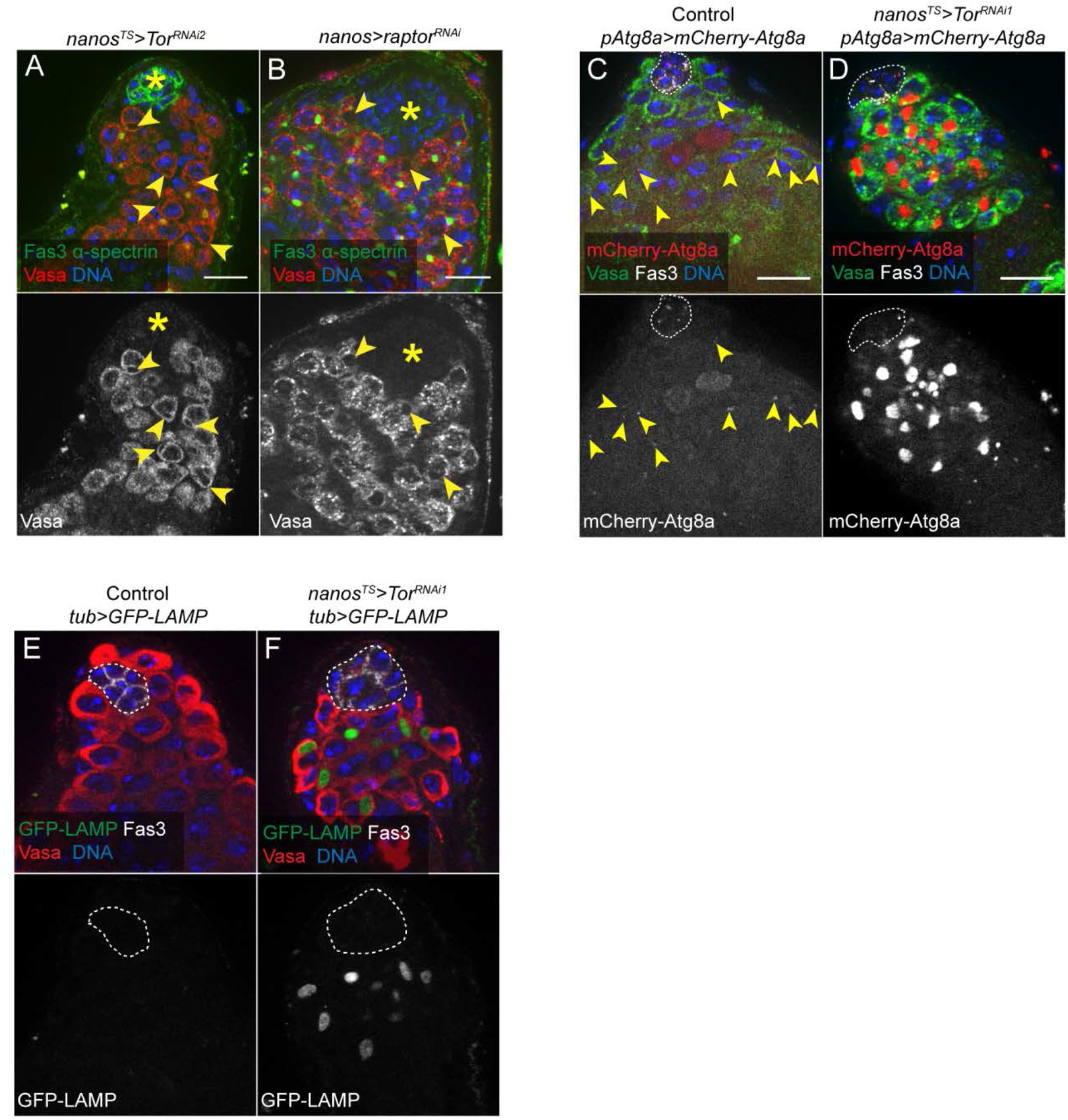
(A-B) Tor-depleted and Raptor-depleted germ cells harbor large cytoplasmic areas devoid of Vasa staining. Images show testis tips from nanos-GAL4/+;UAS-Tor^TRiP-HMS01114^/tub-GAL80^ts^ animals raised at 18°C and shifted to 29°C at the adult stage for 15 days prior to dissection (A) and nanos-GAL4:VP16/UAS-raptor^TRiP--HMS00124^ (B) animals, with Fas3, α-spectrin, Vasa and DNA stainings. Arrowheads point to Vasa-negative structures in the cytoplasm of germ cells. signal large in Tor-depleted germ cells. (C-D) Vasa-negative regions in Tor-depleted germ cells contain Atg8a. Images show testis tips from pAtg8a-mCherry-Atg8a/nanos-GAL4;UAS-TorTRiP-^HMS00904^/tub-GAL80t^s^ animals raised at 18°C and either maintained at 18°C (C) (Control) or shifted to 29°C at the adult stage for 7 days prior to dissection (D), with Vasa, Fas3 and DNA stainings. Arrowheads in (C) point to Atg8a puncta in cyst cells and germ cells in a control testis. (E-F) GFP-LAMP accumulates in Vasa-negative cytoplasmic structures of Tor-depleted germ cells. Images show testis tips from tub-GFP:LAMP1/nanos-GAL4;UAS-TorTRiP-^HMS00904^/tub-GAL80t^s^ animals raised at 18°C and either maintained at 18°C (E) (Control) or shifted to 29°C at the adult stage for 7 days prior to dissection (F), with Vasa, Fas3 and DNA stainings. The hub is indicated with an asterisk in (A-B) and is delimitated with dotted lines in (C-F). For all images, scale bar: 15µm.

**Supplementary Figure 4:**
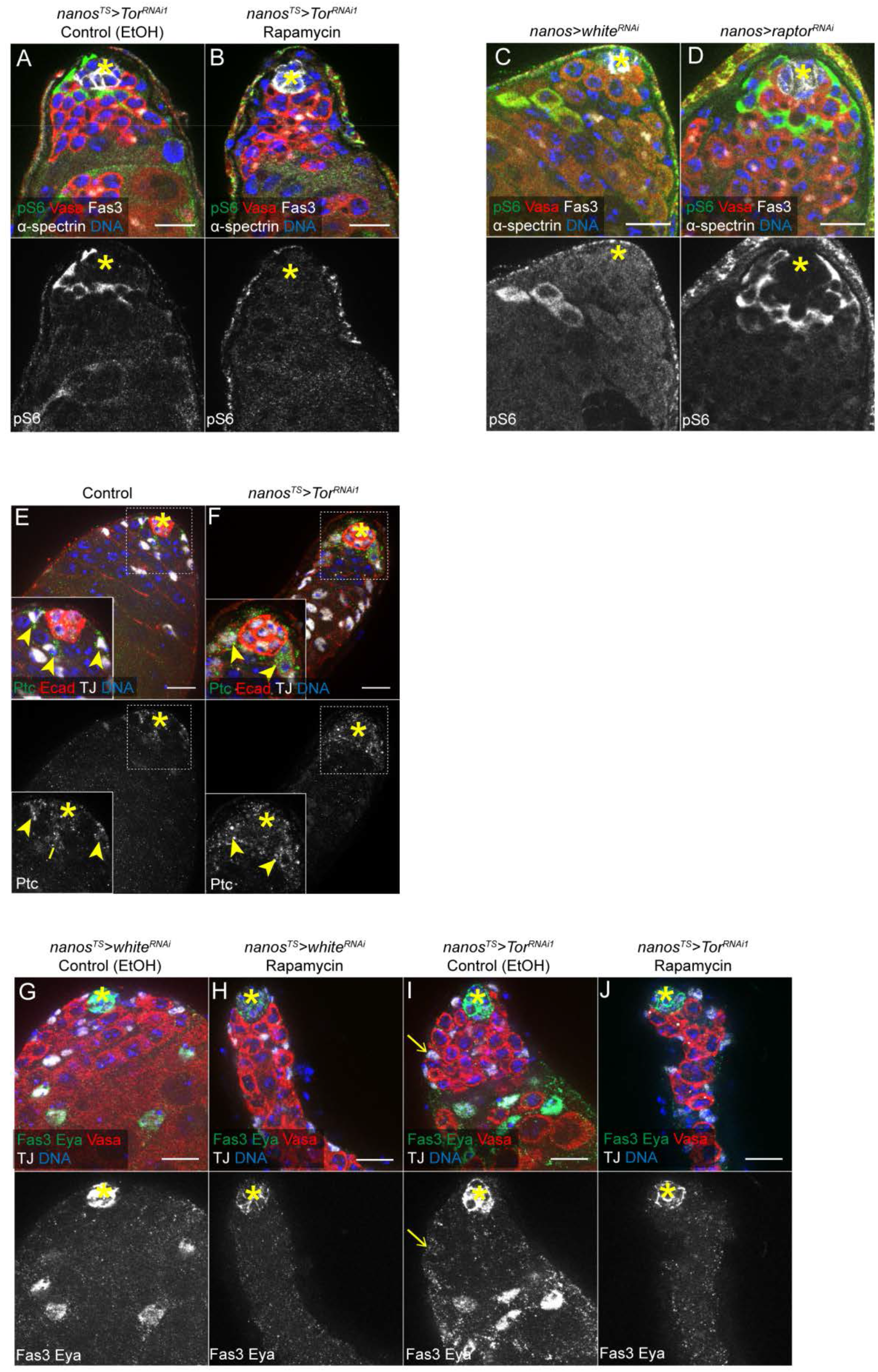
(A-B) Rapamycin treatment prevents the increase of pS6 in cyst cells adjacent to Tor-depleted germ cells. nanos-GAL4/+;UAS-Tor^TRiP-HMS00904^/tub-GAL80^ts^ flies raised at 18°C were shifted to 29°C at the adult stage on food containing either ethanol as a control (A) or rapamycin (4mM in ethanol) (B) for 7 days. Images show testis tips with pS6, Vasa, Fas3, α-spectrin and DNA stainings. (C-D) Depletion of Raptor in germ cells induces an increase of pS6 in neighboring cyst cells. Images show testis tips from nanos-GAL4:VP16/UAS-white^TRiP-HMS00045^ (C) and nanos-GAL4:VP16/UAS-raptor^TRiP--HMS00124^ (D) animals, with pS6, Vasa, Fas3, α-spectrin and DNA stainings. Cyst cells are identified as Vasa-negative and Fas3-negative cells in the testis tip. (E-F) Depletion of Tor in adult germ cells does not affect Patched expression in cyst cells. Images show testis tips from nanos-GAL4/+;UAS-Tor^TRiP-HMS00904^/tub-GAL80^ts^ animals raised at 18°C and either maintained at 18°C (E) or shifted to 29°C at the adult stage for 7 days prior to dissection (F), with Patched (Ptc), E-Cadherin (Ecad), TJ and DNA stainings. Arrowheads point to Ptc-positive and TJ-positive cells in contact to the hub, identified at CySCs. (G-J) Treatment with Rapamycin inhibits Eya expression both in control testes and testes with germline-specific Tor depletion. nanos-GAL4/+;UAS-white^TRiP-HMS00045^/tub-GAL80^ts^ (G-H) and nanos-GAL4/+;UAS-Tor^TRiP-HMS00904^/tub-GAL80^ts^ (I-J) flies raised at 18°C were shifted to 29°C at the adult stage on food containing either ethanol as a control (G and I) or rapamycin (4mM in ethanol) (H and J) for 7 days. Images show testis tips with Fas3, Eya, Vasa, TJ and DNA stainings. Arrows in (I) points to a cyst cell expressing Eya in proximity to the hub. Asterisks denote the hub. All scale bars: 15µm.

